# Stochastic models of polymerization based axonal actin transport

**DOI:** 10.1101/583716

**Authors:** Nilaj Chakrabarty, Peter Jung

**Affiliations:** Department of Physics and Astronomy, Neuroscience Program and Quantitative Biology Institute, Athens, OH 45701, United States of America

**Keywords:** Actin Cytoskeleton, Axonal Transport, Actin Trails

## Abstract

Pulse-chase and radio-labeling studies have shown that actin is transported in bulk along the axon at rates consistent with slow axonal transport. In a recent paper, using a combination of live cell imaging, super resolution microscopy and computational modeling, we proposed that biased polymerization of metastable actin fibers (actin trails) along the axon shaft forms the molecular basis of bulk actin transport. The proposed mechanism is unusual, and can be best described as molecular hitch hiking, where G-actin molecules are intermittently incorporated into actin fibers which grow preferably in anterograde direction giving rise to directed transport, released after the fibers collapse only to be incorporated into another fiber. In this paper, we use our computational model to make additional predictions that can be tested experimentally to further scrutinize our proposed mechanism for bulk actin transport. In the previous paper the caliber of our model axon, the density of the actin nucleation sites to form the metastable actin fibers, the length distribution of the actin trails and their growth rate were adapted to the biologic axons used for measurements. Here we predict how the transport rate will change with axon caliber, density of nucleation sites, nucleation rates and trail lengths. We also discuss why a simple diffusion-based transport mechanism can not explain bulk actin transport.

## 1. Introduction

Actin is a highly conserved protein and one of the key constituents of the neuronal cytoskeleton [1, 2]. Actin plays essential roles in growth cone navigation, cell signaling [3, 4] and has been shown to be critical for synaptic homeostasis [5]. The bulk of axonal actin, like other cytoskeletal proteins, is known to be synthesized in the cell body of the neuron [6], though recent research also suggests that in some cases like migrating fibroblasts, small amounts of actin can be synthesized locally in the axon [7, 8]. From the neuronal cell body, the synthesized actin is supplied to the axon by a process known as axonal transport.

Axonal transport of actin had initially been visualized by injecting radiolabeled amino acids in the optic nerves of guinea pigs and rabbits [9, 10] and by observing the incorporation and movement of these amino acids at around 2 mm/day along the axons. More recently, fluorescent labeling methods have been used to study actin transport which have revealed that actin is transported at rates ranging from 0.2 - 4.0 mm/day and that a significant fraction of the transported actin is present in soluble, monomeric state (reviewed in [11]). However, the mechanism of axonal actin transport still remains unknown. A functional model of other motile axonal proteins like neurofilaments and microtubules that has emerged from live cell imaging and computational studies, is that they are assembled as stable polymers [12, 13] in the cell body, and transported by the molecular motors dynein and kinesin along the axon [14, 15, 16]. Unlike microtubules and neurofilaments, which are known to be stable in the axon, often having turnover time of months [17], actin is known to be highly dynamic, and rapidly exchanging between monomeric and polymeric states. This dynamism is a critical property of actin and is essential for carrying out most of its cellular functions [18, 19, 20]. Hence, a model of active transport via motor proteins in the form of stable, assembled polymers would not be appropriate for actin.

An impediment in visualizing axonal actin has been that GFP tagged actin cannot distinguish between its monomeric and filamentous forms in live cell imaging. The advent of improved imaging methods like super-resolution microscopy and probes like GFP-Utr:CH [21] (Green Fluorescent protein tagged to the calponin homology of utrophin) which bind selectively to filamentous actin (F-actin) has led to the visualization of previously unseen actin-based architectures like actin rings and trails [22, 23]. Actin rings have been shown to form a stable, periodic, circumferential lattice of F-actin and spectrin below the plasmalemmal membrane and plays important structural roles. Actin trails are, in contrast, more dynamic and form a network of deep axonal filaments growing parallel to the axon from focal actin “hotspots” [23]. Actin trails were first visualized in axons of cultured hippocampal neurons [23] and were also observed in *C. elegans* axons via in-vivo imaging [24]. They nucleate on the surface of actin “hotspots” which colocalize with stationary axonal endosomes, observed by barbed-end marking proteins like Mena/Vasp [25] and grow bidirectionally along the axon [23, 26]. Actin trails are formin dependent and treating the axons with the small-molecule formin inhibitor SMIFH2 resulted in a significant decrease of trail nucleation rates, trail lengths and elongation rates. Actin trails polymerize locally in the axon, growing rapidly at rates of nearly 1*µm/s* by consuming monomers from the bulk of the axon and grow to lengths of tens of microns (average 8.8*µm*). Upon depolymerization, the filament releases its monomer content back into the axon. Perhaps the most surprising feature that has consistently emerged from the live imaging data is that actin trails have a small bias (58 % vs 42 % approximately) toward anterograde growth. The dynamism of axonal actin trails and their preference of growing anterogradely suggested that they might play a role in the active transport process.

In a recent paper [26], using live cell imaging, 3D STORM imaging and computational modeling, we demonstrated that actin trails can indeed mediate bulk axonal actin transport at rates corresponding to slow axonal transport. In our model, actin filaments are nucleated locally in the axon, on the surface of stationary hotspots interspersed along the axon. The filaments grow according to established polymerization rates (summarized in [27] and discussed later in section 2.1) and when they depolymerize, their monomer content is deposited back into the axon. Using simulations to track a photoactivated population of actin in the axon, we demonstrated that the bias of actin trail nucleation leads to bulk transport at rates that matched the photoactivation experiments using PAGFP:Utr-CH (PhotoActivable GFP:Utr-CH). In addition, pharmacological disruption of actin trail nucleation and elongation using the small-molecule formin inhibitor SMIFH2 and latrunculin A also led to a collapse of bulk actin transport.

In this paper, we further scrutinize our proposed mechanism for actin transport by making testable predictions of how the bulk actin transport rate depends on axon caliber, basal G-actin concentration, nucleation rates and trail length. We also present a simplified model to approximately predict the transport rate in terms of observable actin trail and axon parameters so that one doesn’t have to execute time-consuming computational simulation to understand their role in actin transport. In this model, instead of considering the biophysics of G-actin/F-actin exchange we consider the population of actin monomers to be in a combination of three different, anterogradely moving, retrogradely moving and freely diffusing states. By formulating a system of partial differential equations to describe the time evolution of a population of actin monomers switching between these three states, we can estimate the bulk transport rate. The results from this simplified model matches closely with the more detailed stochastic model and helps in identifying the important factors affecting the transport rate.

## 2. Methods

### 2.1 Review: stochastic multiscale model of actin trail nulceation, growth and collapse

Actin filament elongation begins with the nucleation step, where actin monomers form a stable trimer complex [28]. The nucleation phase is the rate-limiting step of the actin filament polymerization process since the actin dimer intermediates are highly unstable [29]. Nucleation of actin monomers in cells is facilitated by a host of actin-nucleating proteins which include the Arp 2/3 complex [30], formins [31, 32], and nucleators containing the tandem G-actin binding proteins like Spire [33], cordon-bleu [34] and leiomodin (Lmod) [35]. Actin trails are formin dependent and typically originate from stationary endosomes growing linearly along the axon [23, 26, 36].

To model the actin trail elongation dynamics, we consider a cylindrical axon of radius *r*, in which there is a uniform basal concentration of G-actin. The stationary endosomes along the axon, which serve as anchoring locations for the actin trails, are considered to be point particles in our model, uniformly spaced at a distance of *d*_*h*_ (figure 1). The nucleation of actin trails at each hotspot is considered to be an independent Markov process. If there are no trails growing from an actin hotspot, then either an anterogradely directed or a retrogradely directed trail can be nucleated at the hotspot with rates *r*_*na*_ and *r*_*nr*_ respectively. The nucleation rates of actin trails are derived from the experimentally observed total number of actin trails nucleated (see [23, 26, 21]). If *N*_*a*_ anterograde trails and *N*_*r*_ retrograde trails are observed in an imaging window of length *L*_*w*_ over a time period *T*, the nucleation rates of anterograde and retrograde trails of each hotspot is given by

**Figure 1:**
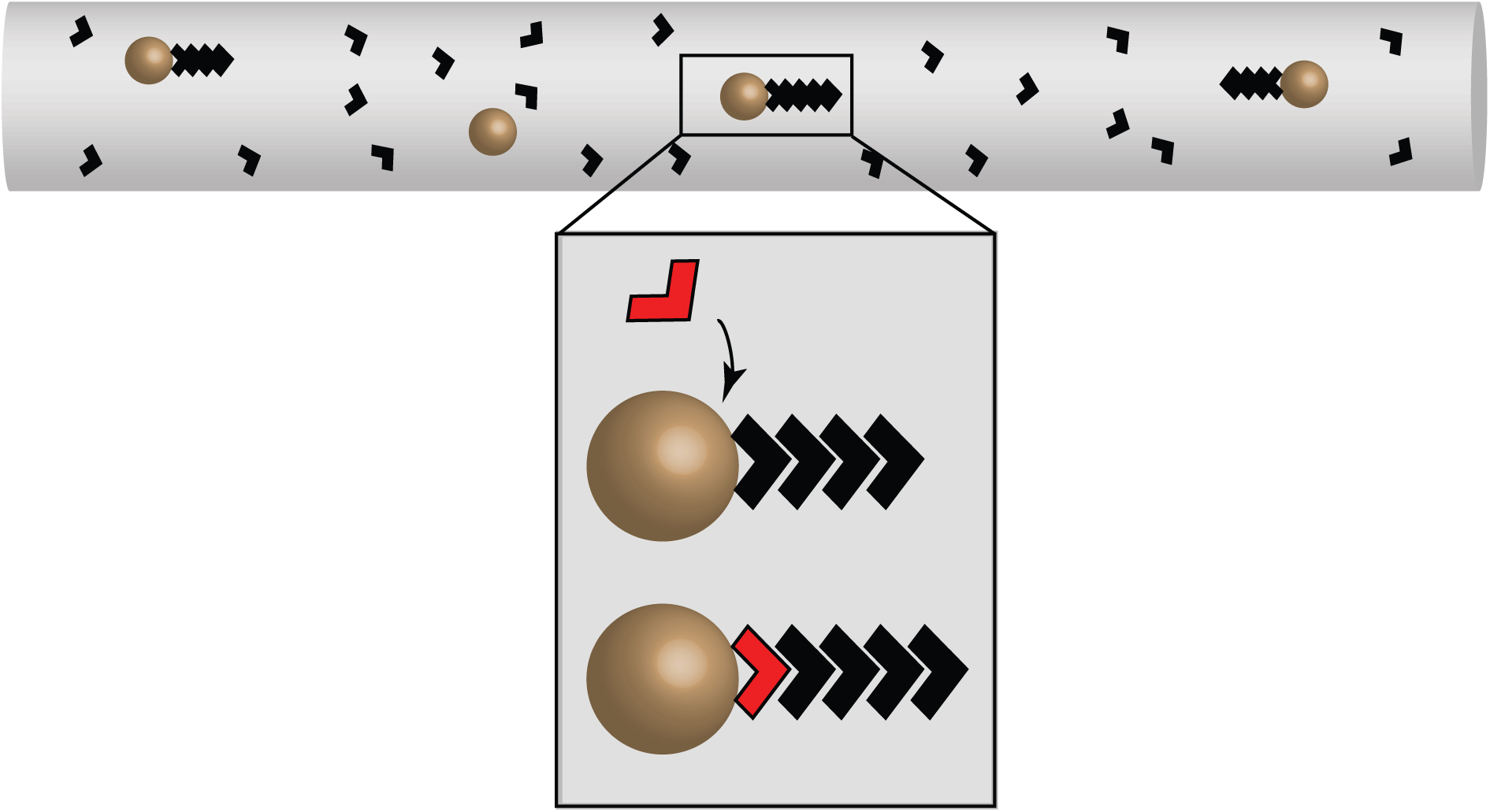
Simulation model of axonal actin trails. Stationary hotspots (brown spheres) are located uniformly along the axon. Actin monomers (black arrowheads) can polymerize as anterograde or retrograde actin trails on the surface of the hotspots. (*Inset)* Progressive movement of monomers on the trail as a new monomer (red) is incorporated at the barbed end of the trail.

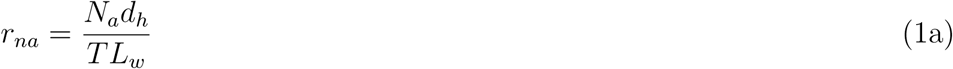

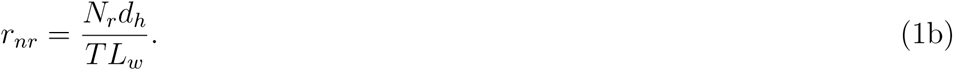

For example, considering the average number of anterograde trails nucleated, *N*_*a*_ = 20.4, and the average number of retrograde trails nucleated, *N*_*r*_ = 14.9, in an imaging window of length *L*_*w*_ = 65 *µm*, where hotspots are spaced at a mean distance of *d*_*h*_ = 3.6 *µm* over a time period *T* = 600*s* (data from [23]), the nucleation rates of anterograde and retrograde trails are *r*_*na*_ = 0.001885*/s* and *r*_*nr*_ = 0.001381*/s* for each hotspot. Due to the relatively low trail nucleation rates of each hotspot, we model the anterograde and retrograde trail elongation as mutually exclusive processes, whereby a hotspot elongating an anterograde trail, for example, cannot simultaneously elongate a retrograde trail. For high nucleation rates this assumption would not be adequate since during the life-time of a trail, the hotspot is not available for other trails to nucleate and there would be a mismatch between the nucleation rates calculated using equation 1 and the observed number of trails.

After a stable multimeric actin nucleus has been formed, the trail grows by incorporating monomers from the G-actin pool of the axon. G-actin can bind either ATP or ADP in its cleft and each species, G-actin-ATP or G-actin-ADP, has distinct association and dissociation rates from the ends of the filament [27, 37]. Due to the high ATP/ADP ratio in the axon [38], most G-actin monomers are ATP bound [39]. Moreover, actin binding proteins like Thymosin-*β*4 and Profilin catalyze the nucleotide exchange of G-actin-ADP to G-actin-ATP [40] which further lowers the fraction of G-actin-ADP pool in the axon. Therefore, we make the simplifying assumption in our model that the unpolymerized pool of G-actin in the axon is only ATP bound. We consider a randomized model of monomer hydrolysis where monomers bound to filaments can hydrolyze independently at a rate of 0.3*/s* to form G-actin-ADP-pi [41] and can release the phosphate group at a rate of 0.0026*/s* to form G-actin-ADP [42]. The rates of G-actin-ATP association and dissociation and that of G-actin-ADP dissociation from the barbed and pointed ends of the filament are summarized in Table 1.

**Table 1:** Reaction rate constants

| Reaction Rate | Description | Value | Reference |
| --- | --- | --- | --- |
| $k_{\text{on,B,T}}$ | Association rate of G-actin-ATP to barbed end | $11.6\mu M^{-1}s^{-1}$ | [37] |
| $k_{\text{off,B,T}}$ | Dissociation rate of G-actin-ATP from barbed end | $1.4 s^{-1}$ | [37] |
| $k_{\text{off,B,D}}$ | Dissociation rate of G-actin-ADP from barbed end | $7.2 s^{-1}$ | [37] |
| $k_{\text{on,P,T}}$ | Association rate of G-actin-ATP to pointed end | $1.3\mu M^{-1}s^{-1}$ | [37] |
| $k_{\text{off,P,T}}$ | Dissociation rate of G-actin-ATP from pointed end | $0.8 s^{-1}$ | [37] |
| $k_{\text{off,P,D}}$ | Dissociation rate of G-actin-ADP from pointed end | $0.3 s^{-1}$ | [37] |
| $k_h$ | Hydrolysis Rate of G-actin-ATP | $0.3 s^{-1}$ | [41] |
| $k_{\text{pi-release}}$ | Rate of pi release from G-actin-ADP-pi | $0.0026 s^{-1}$ | [42] |

The rate of monomer association at the barbed and pointed ends is proportional to the local G-actin concentration at the respective end [43, 37]. The local G-actin concentration is depleted or replenished as monomers are added to or released from the trail. We model the G-actin-ATP (henceforth referred to as G-actin only) concentration along the axon as 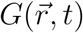 where 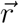 denotes the position vector and *t* time. If an actin trail is growing with its barbed end at position 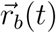 and its pointed end at 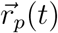, the spatiotemporal change of the G-actin concentration 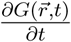 is governed by the equation

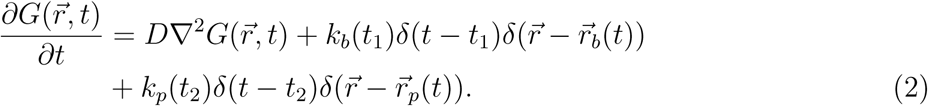

The first term on the right-hand side of equation 2 describes the concentration change due to Fickian diffusion. The diffusion coefficient of actin monomers in cytoplasmic medium is known to be around 4.5 − 7 *µm*^2^*/s* [44] and is taken to be 6 *µm*^2^*/s* for our model. The second term describes the instantaneous concentration change when a monomer is added to or released from the barbed end of the trail located at 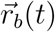 at time *t* = *t*_1_. The pre-factor *k*_*b*_(*t*) is determined stochastically at each step by choosing from the set of possible reactions at the barbed end of a filament using the Gillespie algorithm [45]. If a G-actin monomer is added to the barbed end of the trail at time *t*_1_, then there will be an instantaneous depletion of G-actin at that location 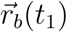, and *k*_*b*_(*t*_1_) = −1. If a monomer is released from the barbed end of the trail at time *t*_1_ then *k*_*b*_(*t*_1_) = +1. Similarly the third term describes the instantaneous concentration change due to a monomer being added to or released from the pointed end of the filament. Note: the units of length and time are *µ*m and seconds, respectively. To convert the units of concentration from 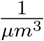 to *µ*M, a conversion factor of 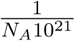 has to be multiplied to *k*_*b*_ and *k*_*p*_, where *N*_*A*_ is the Avogadro’s number.

In our model, the nucleotide bound to each actin monomer in the trail is tracked over time since G-actin-ADP has different dissociation rates from each end of the trail. However, since we consider the monomer pool to be composed of only G-actin-ATP, we do not track the G-actin-ADP in the axon pool over time.

The diameter of the axons (ranging from 140 − 200 *nm*) [23] is very small compared to its length and the concentration profile is relatively constant along the radial direction. Therefore, we assume the G-actin-ATP concentration to be constant along the radial direction of the axon and we integrate equation 2 along the radial direction to obtain the associated one-dimensional equation (see Appendix resultant one-dimensional equation governing the G-actin-ATP concentration along the length of the axon is therefore given by

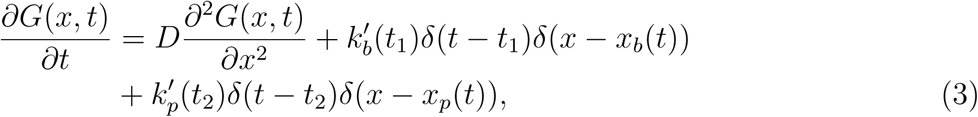

where 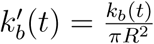 and 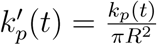 and *R* is the axon radius.

We solve equation 3 using periodic boundary conditions,

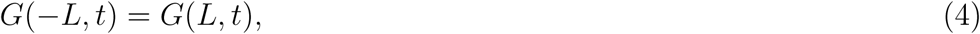

which ensures a flux balance in the axon such that the flux of the G-actin-ATP monomers at the retrograde end of the axon matches that at the anterograde end.

Since the trails grow with their barbed ends attached to stationary hotspots, the position of the barbed end of each trail, *x*_*b*_(*t*), is taken to be constant over time, while the position of the pointed end, *x*_*p*_(*t*), changes over time as new monomers are added to or dissociates from either ends of the ends (figure 1). Since the barbed end of the trail stays in position, the constituent monomers of the trail are progressively moved forward as new monomers are added at the barbed end (figure 1, inset). We model the collapse of actin trails as an instantaneous process, where once an actin trail reaches its collapse length, it instantly deposits its monomer content back into the axon. The collapse lengths of the trails are drawn from the frequency distribution of the length of trails observed in the imaging experiments using GFP:Utr-CH [23] and plotted in figure 2.

**Figure 2:**
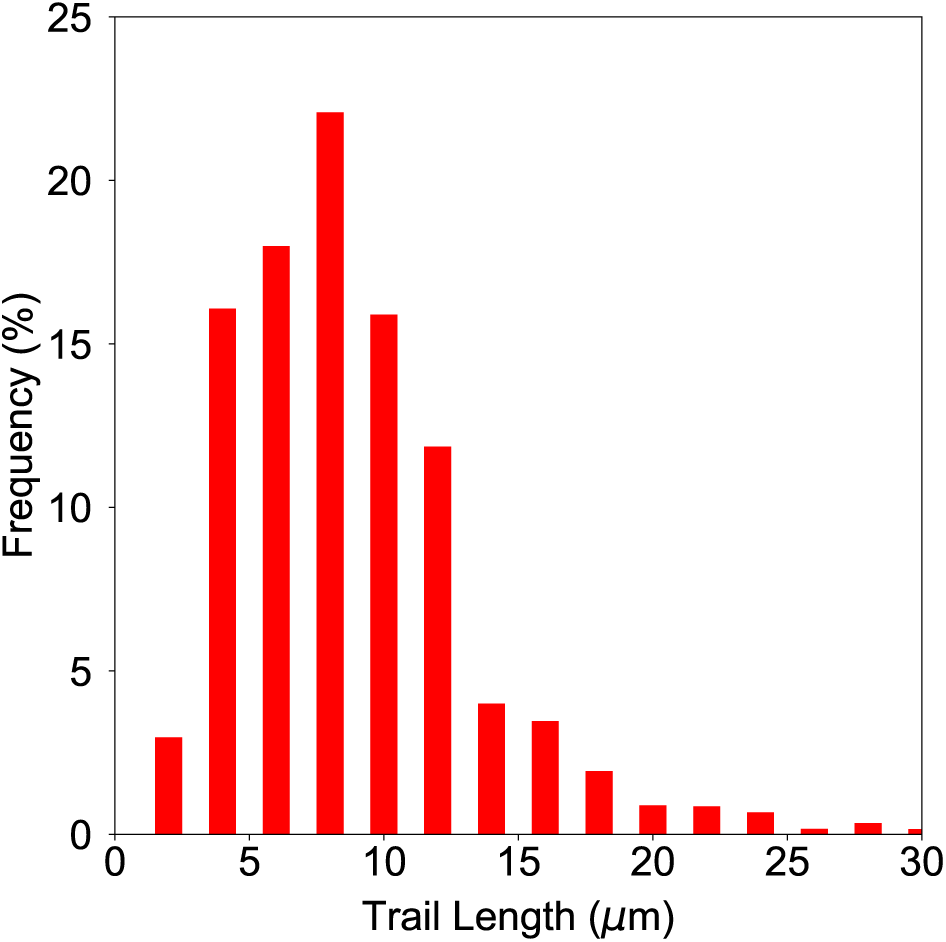
Length distribution of actin trails incorporated in the simulations. Reproduced using data from [23].

Due to the higher nucleation rates of anterogradely directed actin trails and since attached monomers are processively pushed forward, we expect a net anterograde transport of actin. Actin monomers diffuse passively in the axon and are occasionally translocated when incorporated into growing trails. To quantify the rate of transport, such that it can be directly correlated with similar experiments, we computationally replicate a photoactivation experiment.

At the start of the simulations (taken to be *t* = −50 *s*), actin is present only in its monomeric form as G-actin-ATP. At *t* = 0, after monomeric and filamentous actin reaches a steady state of exchange, both G-actin and F-actin, in a central zone of the axon, 15 *µm* length (in accordance with usual photoactivation protocol), is photoactivated. At *t* = 0, immediately after photoactivation, the distribution of fluorescently active G-actin is given by

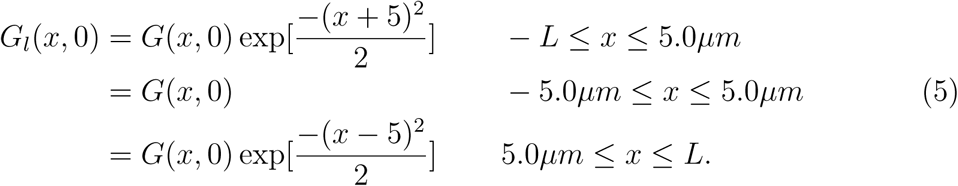

In addition, the monomers on a trail located in this zone are also photoactivated. The dynamics of both the fluorescently active and inactive G-actin is modeled using equation 3, while the trail nucleation and growth is simulated using the Gillespie algorithm [45] with rates *r*_*na*_ and *r*_*nr*_ and the monomer incorporation rates summarized in Table 1. While we used periodic boundary conditions to maintain a flux balance of fluorescently inactive G-actin, for the fluorescently active G-actin, we use Dirichlet boundary conditions

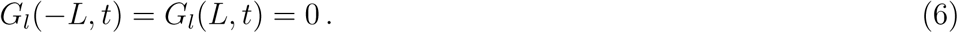

We track the photoactivated actin until a significant amount of it (more than 1%) is lost from the ends of the axon. Hence this particular boundary condition does not affect the results of our simulations. To quantify the transport rate, we calculate the position of the center of fluorescence by measuring its position over time using the equation

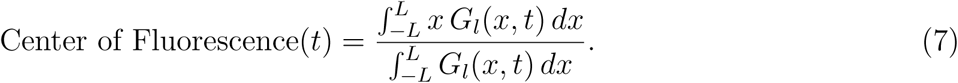

The center is then averaged over hundreds of independent simulation runs and fitted as a function of time to a straight line. The transport velocity is given by the slope of this line.

### 2.2 A stochastic 3-state model of actin transport in presence of diffusion

As seen from the detailed stochastic model of actin trail nucleation and elongation in the axon, an actin monomer exists in three different states, either moving anterogradely (when incorporated in an anterograde trail), moving retrogradely (when incorporated in a retrograde trail) or freely diffusing. An alternative way to model actin transport is to consider the transition dynamics of actin monomers between these three different kinetic states, which we describe as the 3-state model.

The three states are termed *a, r* and *d*. When an actin monomer is in either the anterograde *a* or retrograde state *r*, i.e. when it is incorporated into an anterogradely or retrogradely growing trail, it moves with the trail elongation velocities *v*_*a*_ or *v*_*r*_. A monomer in the anterograde *a* or the retrograde state *r* can transition to the freely diffusing state *d* with rates *γ*_*ad*_ and *γ*_*rd*_ which are associated with the trail disintegration rates. When a monomer is in the freely diffusing state *d*, it can transit into the anterograde state or retrograde state with rates *γ*_*da*_ or *γ*_*dr*_ which are associated with the trail nucleation rates.

The transition scheme is summarized in figure 3. To link the kinetics of actin monomers to spatiotemporal distributions *C*_*a*_(*x, t*), *C*_*r*_(*x, t*) and *C*_*d*_(*x, t*) of large ensembles of monomers in the anterogradely moving, retrogradely moving and freely diffusing states, we consider the system of partial differential equations

**Figure 3:**
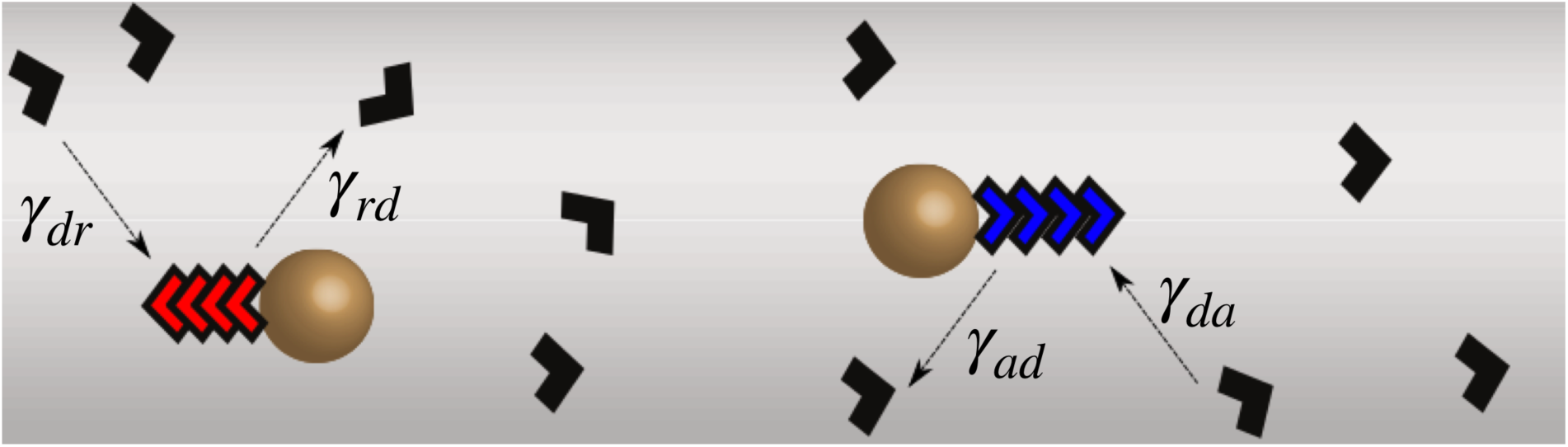
Stochastic 3-state model of actin transport. In this model, actin monomers exist in three different kinetic states, anterograde (blue arrowheads), retrograde (red arrowheads) and freely diffusing (black arrowheads). A monomer can transition from the freely diffusing state to either the anterograde or retrograde states with rates *γ*_*dr*_ or *γ*_*dr*_ respectively. While in either the anterograde or retrograde states, the monomers can also transition to the freely diffusive state with rates *γ*_*ad*_ or *γ*_*rd*_ respectively.

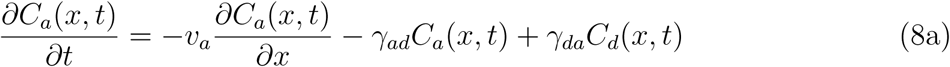

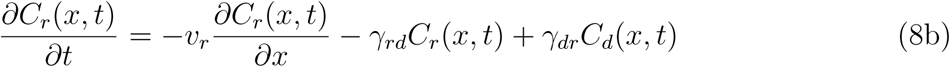

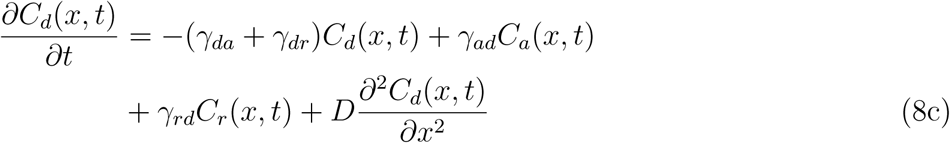

The steady state transport velocity 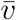 is solved from this system of equations and expressed in terms of observable features of the trails like trail nucleation rates, trail elongation velocity and mean length of actin trails. In homogenous steady state, i.e. 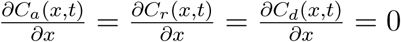 and 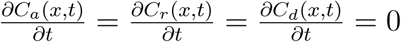, we have

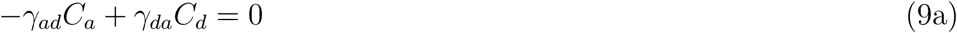

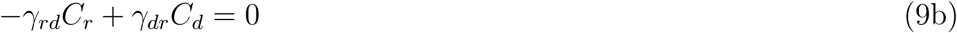

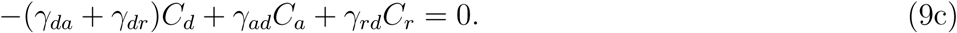

*C*_*a*_, *C*_*r*_ and *C*_*d*_ are the amounts of actin in each kinetic state when the system has reached steady state. The total amount of actin in all the states is given by *C* = *C*_*a*_ + *C*_*r*_ + *C*_*d*_ and is equivalent to the basal G-actin concentration due to the conservation of mass. From equations 9a and 9b we have

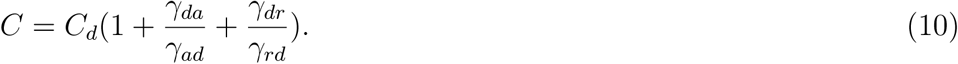

Let 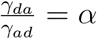 and 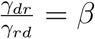. The fraction of actin in the different kinetic states are

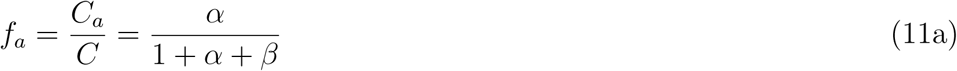

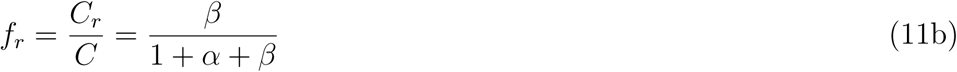

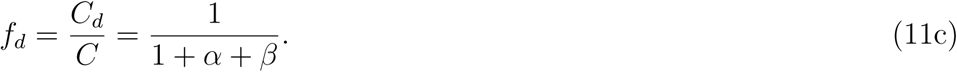

The steady state transport velocity is given by

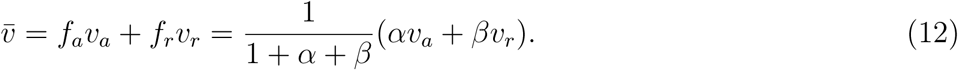

Since the anterograde and retrograde trail elongation velocities are equal in magnitude and directed opposite to each other, *v*_*a*_ = *-v*_*r*_ = *v*. Hence,

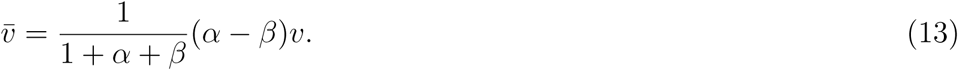

We approximate the trail elongation velocity, *v* as a linear function of the global G-actin concentration, *C* and dependent on the monomer association and dissociation rates at the barbed end of the trail (discussed in detail in section 3.2 and demonstrated in figure 8,a).

**Figure 8:**
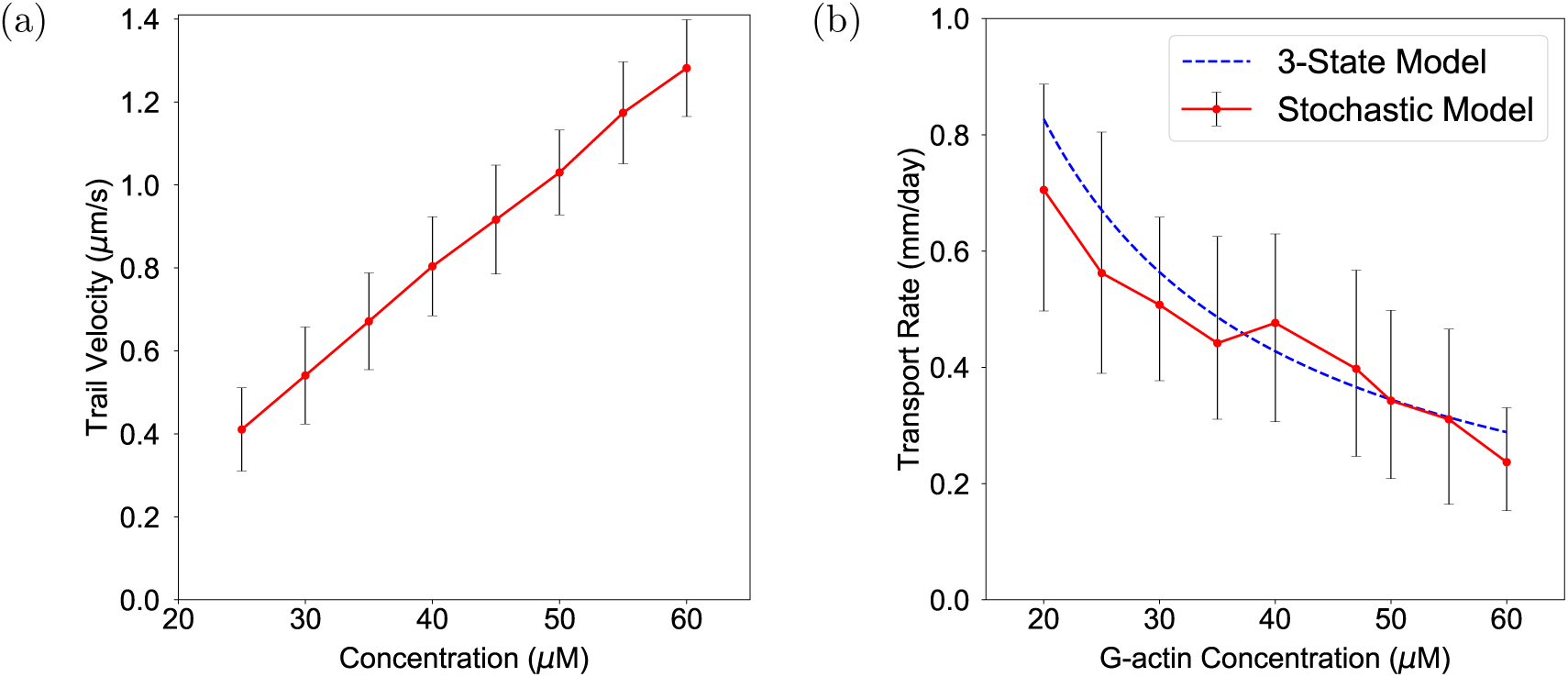
G-actin concentration

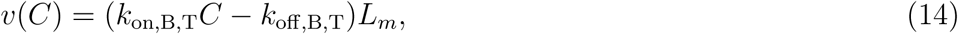

where *L*_*m*_ = 2.7 *nm* is the half-length of the actin monomer (filament length increment due to addition of a monomer).

Next we derive the expressions for *α* and *β*, in terms of observable trail parameters so that the transport velocity can be obtained analytically. To do this, we need to estimate the mean fraction of monomers in each of the three kinetic states. This can be calculated from the nucleation rate of the hotspots, the length of the trails and the elongation velocity of the trails.

We consider the stochastic model of actin trail nucleation, growth and collapse introduced in section 2.1. The cylindrical axon of radius *r* and length *L* has actin hotspots uniformly spaced at a distance of *d*_*h*_. The anterograde and retrograde trail nucleation rates (*r*_*na*_, *r*_*nr*_) are found from equations 1a and 1b (by replacing the imaging window length, *L*_*w*_ with the axon length, *L*_axon_). As per the 3-state model, we assume that once a trail has been nucleated it grows with a uniform elongation velocity *v* before it collapses and deposits its monomer content back in the axon. The elongation velocity does change over the lifetime of a trail due to monomer depletion from the axon (discussed in detail in section 3.1) and therefore the trail velocity decreases with increasing trail length. However, by taking *v* as the average trail velocity, we are assuming this effect is averaged out.

First we deduce the ratio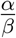. Since 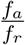 is the ratio of the monomers in anterograde state to the monomers in retrograde state, and the rate of collapse of trails are identical, this ratio must be equal to the ratio of trail nucleation rates. Hence,

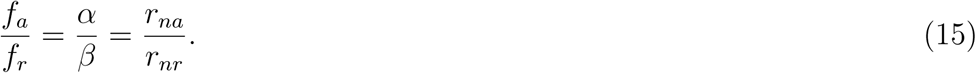

Next we determine the fraction of monomers in the diffusive state, which is the fraction of monomers not bound on trails. We calculate the probability that a hotspot is growing a trail of length *L*_*t*_. The hotspot is like an excitable system. After it spawns a trail and the trail collapses, the average time to generate a new trail is

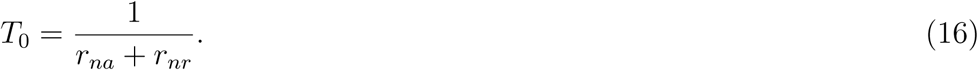

The hotspot is excitable during this time interval. Once a trail of length *L*_*t*_ is nucleated, it grows for a time 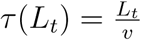 where *v* is the elongation velocity of the trail. During this time, the hotspot is refractory and cannot generate a new trail. The fraction of time the hotspot is refractory is then 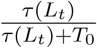. Hence, the probability that a hotspot is growing a trail of length *L*_*t*_ is

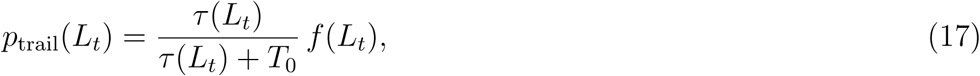

where *f* (*L*_*t*_) is the frequency distribution of trail lengths. To estimate the fraction of monomers bound as trails, we calculate the mean length of trails growing along the axon. Since the trail nucleation process is an independent event, the mean number of hotspots growing a trail of length *L*_*t*_ is

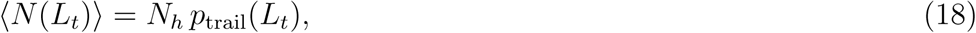

where *N*_*h*_ is the total number of hotspots along the axon. The hotspots that grow a trail of length *L*_*t*_ nucleate randomly and independent of each other. Hence, the mean trail length considering only hotspots that are growing trails of length *L*_*t*_ is the mean number of hotspots times half the maximum length. I.e.,

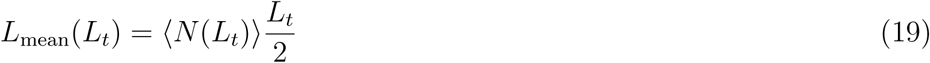

Therefore, the mean total length of actin trails growing in the axon at any time is obtained by integrating over the length distribution

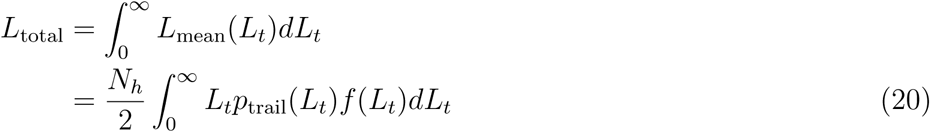

The mean total number of bound monomers in the steady state is 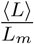. Therefore,

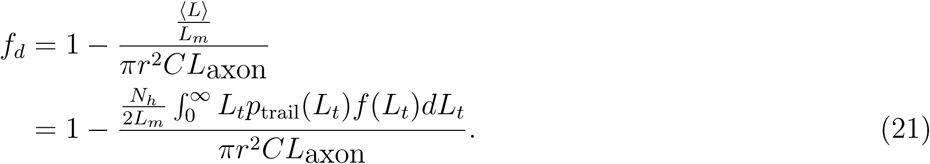

From equations 11c and 21

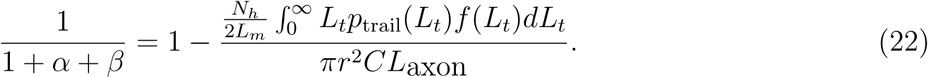

The transport rate 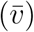 can be obtained in terms of observable trail parameters by simultaneously solving for *α* and *β* using equations 15 and 22

## 3. Results

### 3.1. Diffusion driven transport

To underscore the importance of an active transport process to convey actin from its synthesis sites in the cell body of the neuron, we first consider passive transport driven by a concentration gradient only. We assume that the concentration along the radial direction is constant and focus on the diffusion along the length of the axon. For a source located at *x* = 0 producing a constant flux of *j*_in_, the diffusion equation along the axon can be written as

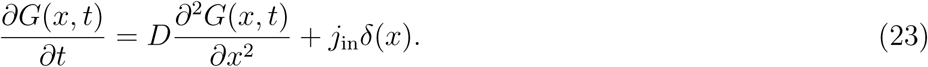

We consider the entire domain to range from *−L* to *L* and the ends of this domain to be held at constant concentration *G*_0_. Half of the flux enters the axonal domain (ranging from 0 to *L*) and the steady state solution in this domain is given by

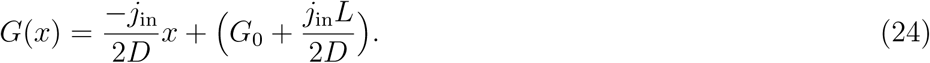

The diffusive flux is given by

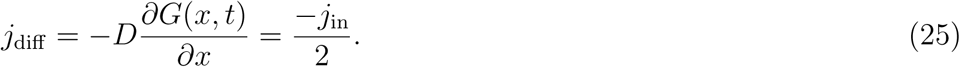

If the concentration gradient in the axon is driving a bulk transport at velocity *v*, the constant flux of the source can be found by equating it to the diffusive flux, i.e.,

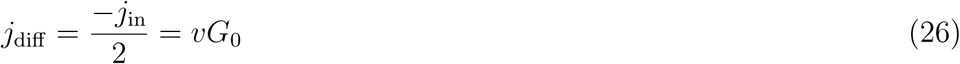

For a diffusion coefficient of *D* = 6 *µm*^2^*/s*, a transport velocity of *v* = 0.4 mm/day = 0.0046 *µm/s* and the basal G-actin concentration of 47 *µM*, the resultant concentration gradient along the axon is given by

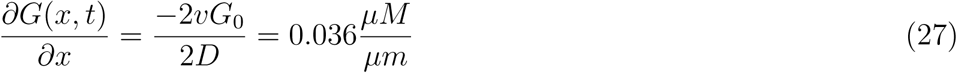

For a concentration gradient of this magnitude, across axons which are hundreds of micron or even a few millimeter long, this predicts a concentration drop of almost 20 to 100 %. No such persistent concentration gradient of G-actin is generally observed along the axon [46]. Since the velocity of trail elongation is directly proportional to the local G-actin concentration, this gradient also predicts that the observed trail velocity would steadily decrease along the axon. However, the trail elongation velocity is not correlated to its nucleation position along the axon [23, 24]. Hence, even if some transport occurs due to the concentration gradient when monomers pour in from the cell body into the axon, this cannot significantly account for the bulk transport of actin.

### 3.2. Actin trail dynamics and transport rate measurement

To analyze the dynamics of a growing actin trail we simulate a single actin trail growing from a hotspot situtated at *x*_*b*_ = 0 (the center of the axon). In figure 4, we plot the G-actin concentration at the hotspot location (*G*(0, *t*), figure 4 upper panel) and the length of the actin trail (figure 4, lower panel). The concentration is initially at its basal value of 47.0 *µM* (explained in the next paragraph). As the trail is nucleated at *t* = 26.8 *s*, it starts incorporating monomers from the axonal pool and the concentration at *x*_*b*_ = 0 starts declining. The declining concentration leads to the trail growth velocity slowing over time, as seen from the decreasing slope of the length of the trail with respect to time. The trail collapses as it reaches a length of 8.8 *µm* after 8.6 *s* and deposits its monomer content along the axon at which time the concentration at the hotspot recovers to its basal value.

**Figure 4:**
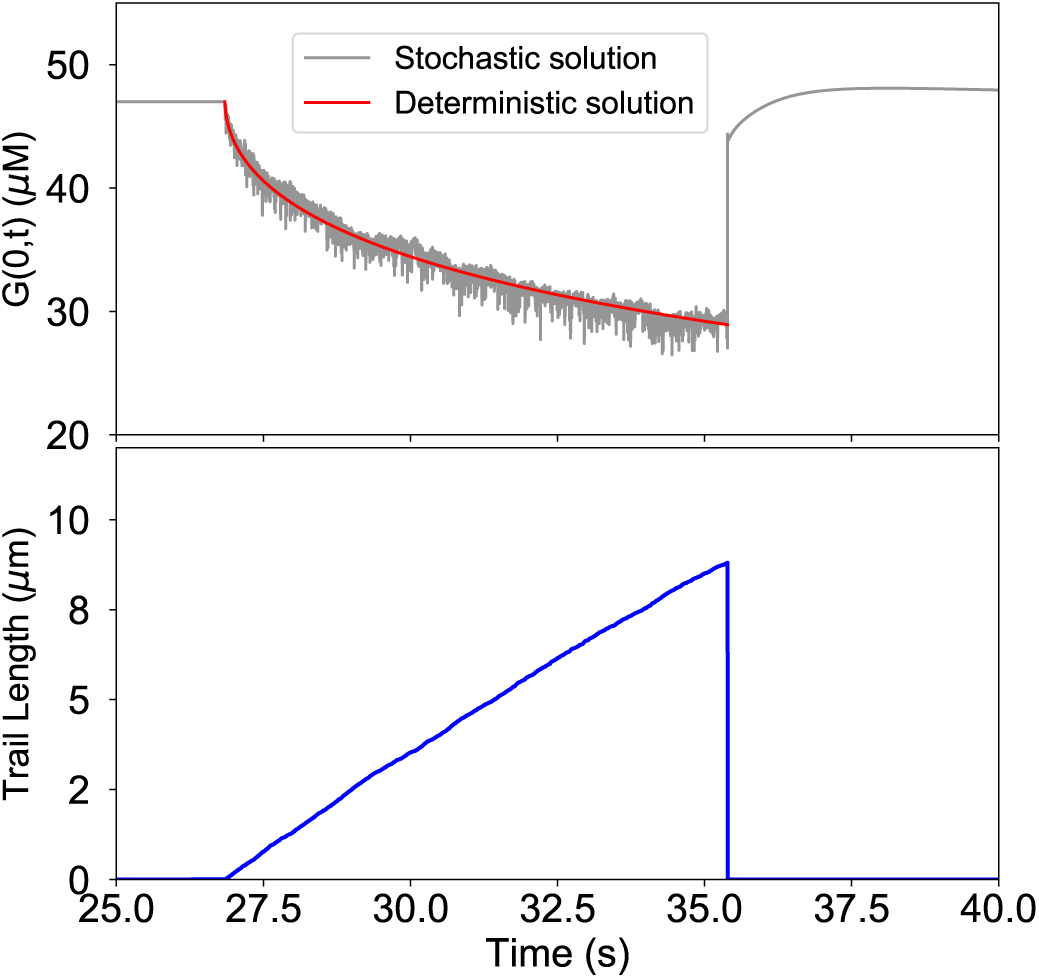
Decline of axonal G-actin concentration during trail growth. The G-actin concentration at the location of a hotspot at *x*_*b*_ = 0 (upper panel) and the length of the trail (lower panel) are plotted as a function of time. The concentration of G-actin at the barbed end is initially at its basal value of 47 *µM*. An actin trail is nucleated from this hotspot at *t* = 26.8*s*. The G-actin concentration in the axon is governed by equation 3. The declining phase of the concentration at the barbed end can also be approximated by equation 30 (upper panel, red line). When the trail reaches a length of 8.8 *µm* after 8.6 *s*, it collapses and the monomer content of the trail is deposited in the axon and the concentration recovers to its basal value.

In an infinite linear system, with initial uniform concentration *C*_0_, if an amount *M* is removed instantaneously from *x* = 0, the concentration at (*x, t*) is obtained by solving the one-dimensional diffusion equation and is given by

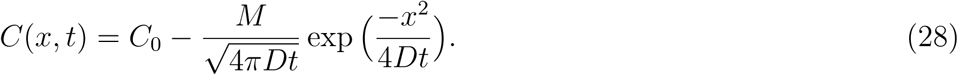

If the diffusing substance is removed continuously at a rate *q*(*t*′) from *x* = 0, the resultant solution is obtained by integrating equation 28 till time *t* and is given by

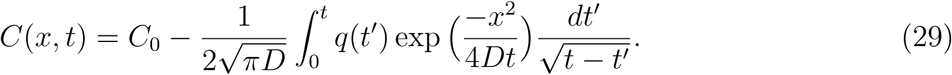

The declining phase of the G-actin concentration due to monomers being absorbed at the barbed end at *x*_*b*_ = 0 can hence be approximated by the equation

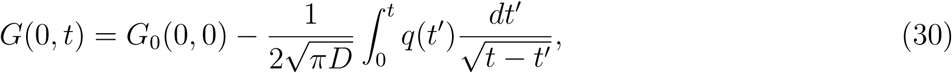

where *G*_0_(0, 0) is the basal concentration of G-actin at the hotspot location *x*_*b*_ = 0 and *q*(*t*′) = *k*_on,B,T_ *G*(0, *t*′) is a sink term dependent on the instantaneous concentration of actin, *G*(0, *t*) and the rate constant of incorporating monomers at the barbed end of the trail, *k*_on,B,T_. The resultant equation 30 is a linear Volterra equation of the second kind [47]. This equation neglects the incorporation of the monomers at the pointed end of the trail which accounts for approximately 8 % of the monomers but provides a reasonable estimate of the concentration at the barbed end of the trail. The numerical solution of equation 30 is compared with the stochastic result in figure 4 (upper panel, blue line) [26].

Actin trail growth velocity is significantly affected by the basal concentration of G-actin in the axon. The G-actin concentration in the axon is estimated by matching the mean trail growth velocity to the experimentally observed value of 1 *µm/s* [23] (also discussed in section 3.3.3). At 47 *µM* G-actin concentration, the mean trail elongation velocity is 1 *µm/s* which matches the experimentally observed trail elongation rate.

We simulate axonal actin trails with anterograde and retrograde nucleation rates of *r*_*na*_ = 0.001885*/s* and *r*_*nr*_ = 0.001381*/s* for each hotspot in an axon of diameter 170 *nm*. Following the photoactivation procedure described in the methods section, in figure 5, we plot the center of the photoactivated zone, averaged over 1000 runs and fit it to a straight line to obtain the transport rate. The transport rate is found to be 0.0046 *µm/s* or 0.39 mm/day which corresponds to slow axonal transport rates.

**Figure 5:**
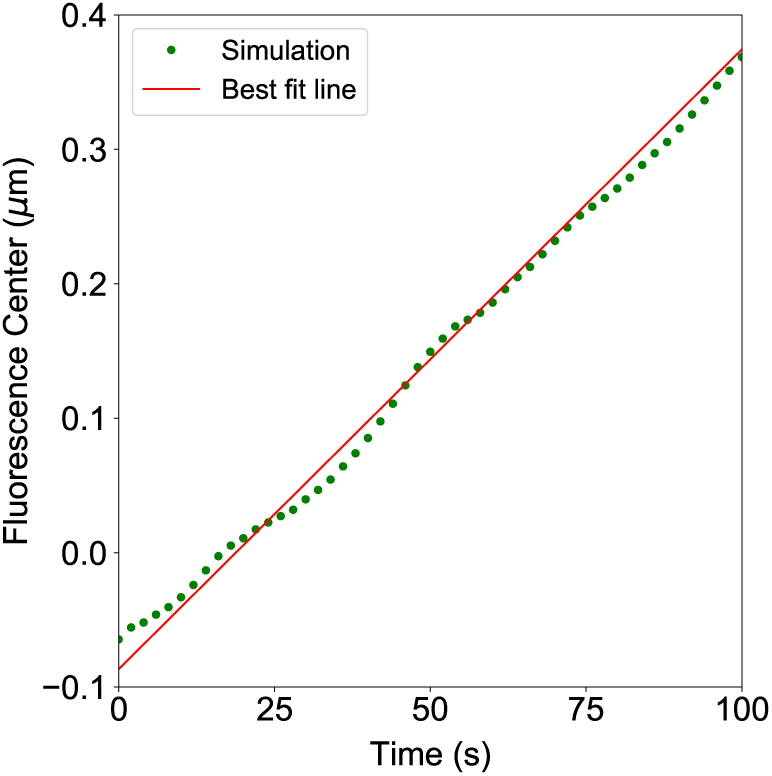
Transport rate determination from a photoactivation simulation. The anterograde and retrograde nucleation rates are *r*_*na*_ = 0.001885*/s* and *r*_*nr*_ = 0.001381*/s*, the basal G-actin concentration is 47*µM*, the axon radius is *r* = 85 *nm*. The simulation begins at *t* = −50 *s* and the fluorescently tagged monomers are activated at *t* = 0 *s*. The center of fluorescence is averaged over 1000 independent runs and fitted to a straight line. The slope of the best fit line is 0.0046 *µm/s* (0.39 mm/day).

Plugging in these parameters for the trail nucleation rates, the basal G-actin concentration and the axon radius in equations 15 and 22 we can solve for *α* and *β* of the 3-state model and from equation 13, the steady state transport velocity is calculated as 0.36 mm/day which is close to the rate measured using the detailed stochastic model (0.39 mm/day) and well within error estimates (demonstrated in the next section and discussed in detail in Appendix B).

### 3.3. Effect of axonal and trail parameters on the transport rate

In this section we discuss the dependence of the bulk actin transport rate on various axonal and actin trail parameters. The important parameters in our model are the nucleation rate of actin trails, the radius of the axon, the basal concentration of G-actin and the length of actin trails. In each simulation, the trail elongation velocity is found by averaging over 1000 trails and the transport rate is found using the photoactivation algorithm described in the last section averaged over 1000 runs. The distribution of the trail velocities for each data point is characterized by the standard deviation of elongation velocities. The distribution of the transport velocities is quantified by linearly fitting the mean *±* 95% confidence interval of the center of fluorescence over time (described in detail in Appendix A).

#### 3.3.1. Trail nucleation rates

To study the dependence of the transport rate on the nucleation rate, we introduce a nucleation prefactor parameter (*k*_*p*_) and express the anterograde and retrograde trail nucleation rates as *k*_*p*_× observed rates. Therefore, *r*_*na*_ = *k*_*p*_ × 0.001885*/s* and *r*_*nr*_ = *k*_*p*_ × 0.001381*/s*. This ensures that as the nucleation prefactor is varied, the bias 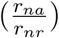 is constant, but the total number of trails change. Physiologically, this represents a situation where the filament-nucleating factors like formin are increased. Such nucleating factors are known to affect trail nucleation significantly while having limited effect on trail elongation rates [23].

The effect of changing the nucleation prefactor on the trail elongation velocity and the transport rate is shown in figure 6. As the total number of trails increase, there is a slight decrease in the average trail elongation velocity, as seen in figure 6(a), due to the effect of monomer depletion, since the constant basal concentration of G-actin (47 *µM)* is supplying an increasing number of trails. However, even though trails slow down slightly with increasing number of trails, there is an overall increase of the transport rate (figure 6(b) red solid line) since there are more monomers being transported in the anterograde direction, even for the same bias. In the 3-state model the trail elongation velocity does not change with the nucleation factor since the effect of monomer depletion is not accounted for. The steady state fraction of monomers in both the anterograde and retrograde state (*f*_*a*_, *f*_*r*_) increases with increasing *k*_*p*_ since the number of trails increases. Hence even though the bias 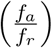 remains same, there is an increase in the transport velocity, 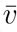, due to an increase in their absolute difference as shown in figure 6(b) (blue dotted line).

**Figure 6:**
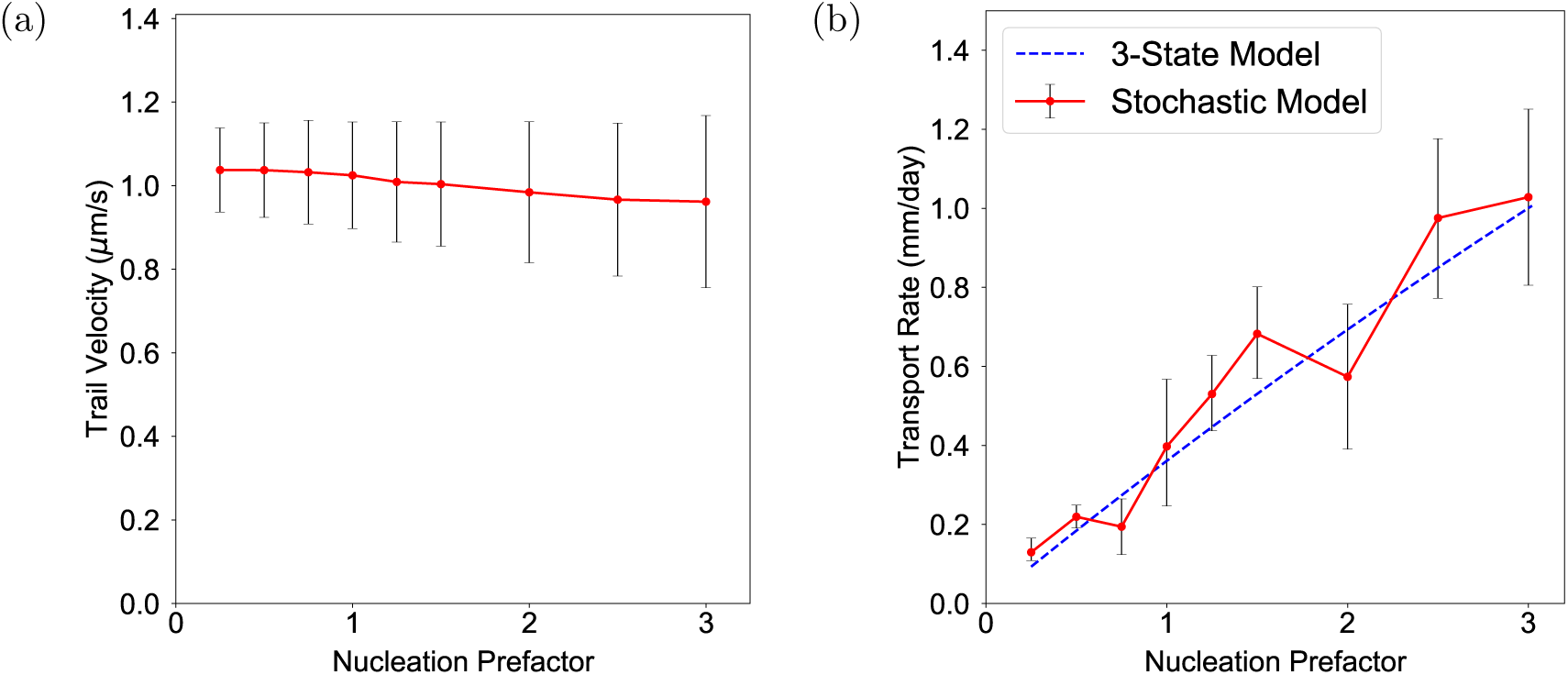
Trail elongation velocity and slow transport rate as a function of the total number of trails. The total number of trails increase as the nucleation prefactor parameter is increased, though the bias of the trails is constant. (a) The trail elongation velocity is averaged over 1000 trails of all length for each nucleation prefactor value. (b) The transport rate is calculated for each nucleation prefactor value by performing a photoactivation simulation and averaging over 1000 runs (similar to figure 5) and from the 3-state model.

#### 3.3.2. Axonal radius

Next, we study the effect of axonal radius on the trail elongation velocity and the transport rate. We vary the axonal radius from 70 nm to 100 nm, the physiological limits that are observed in the imaging experiments. Experimentally, this can be tested by measuring the transport rate in various developmental stages of the animal which is characterized by axons of increasing caliber.

With an increasing radius, there is an increase in the trail elongation velocity (figure 7(a)) since for the same G-actin concentration, the total number monomers to supply the growing actin trails is increasing. However, for the same number of trails, due to an increase in the monomer content in the axon, a lesser fraction of the total number of monomers are now transported via trails. The overall effect is a decrease in the transport rate with increasing axonal radius (figure 7(b), red solid line). Similarly, in the 3-state model, with an increasing radius, there is an increase in the steady state fraction of monomers in the freely diffusing state (*f*_*d*_). The fraction of monomers in both anterograde and retrograde states (*f*_*a*_, *f*_*r*_) therefore decreases. Though the velocity of the trails is assumed to be constant in the 3-state model, this leads to a decline in the transport rate (figure 7(b), blue dotted line). Physiologically this suggests that actin transport in axons of bigger caliber which have a higher actin requirement have to be supplemented either by higher trail nucleation rates or by other factors like stabilizing the trails themselves (discussed in section 3.3.4).

**Figure 7:**
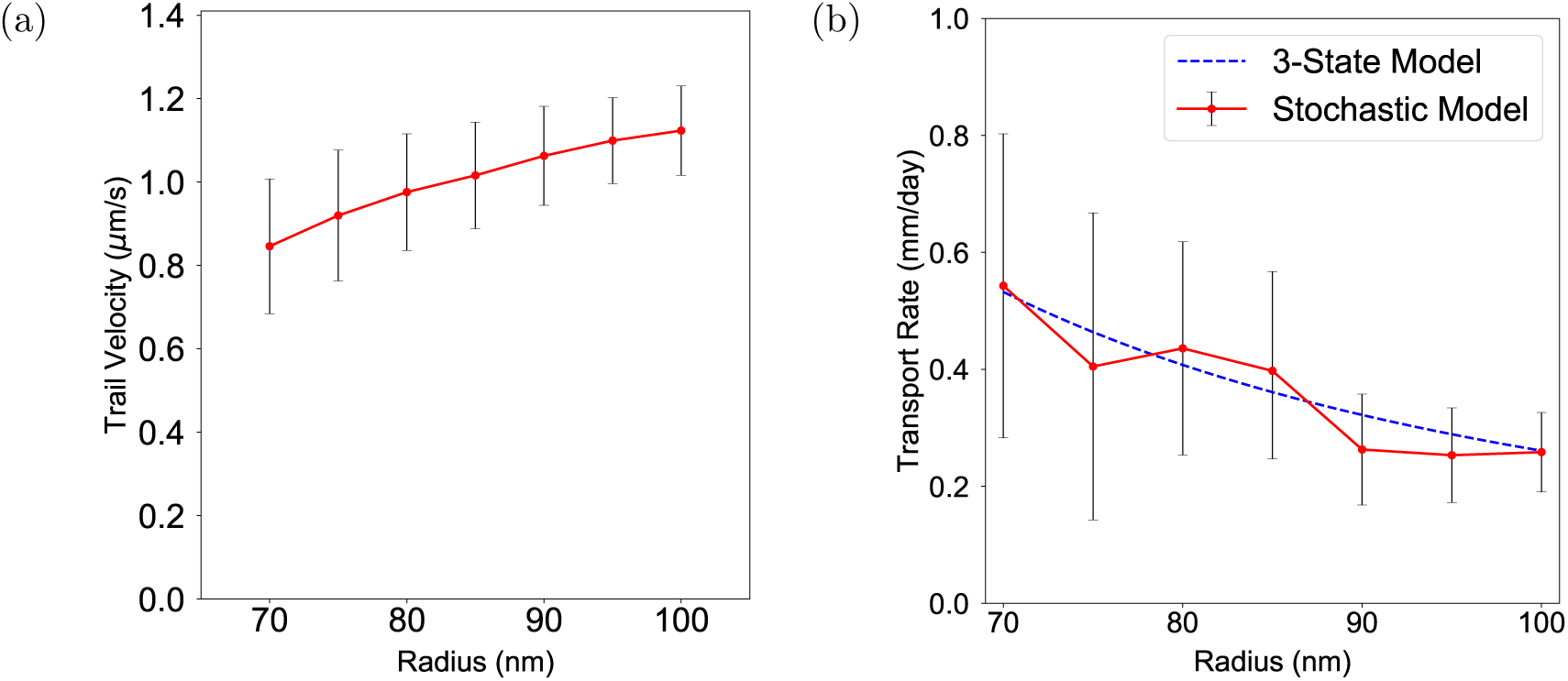
Trail elongation velocity and slow transport rate as a function of the axonal radius. (a) The trail elongation velocity is averaged over 1000 trails of all length for each radius. (b) The transport rate is calculated for each radius by performing a photoactivation simulation and averaging over 1000 runs.

#### 3.3.3. Basal G-actin concentration

Actin trails elongate by absorbing monomers from the bulk of the axon and the rate of monomer association is directly proportional to the bulk G-actin concentration. With increasing G-actin concentration, the trail elongation velocity increases linearly as seen in figure 8,a. Here, the trail elongation velocity is averaged across trails of all length.

As seen in figure 8 b, the effect of increasing concentration on the transport rate is qualitatively similar to the effect of increasing radius discussed in the last paragraph. Even though for an increasing G-actin concentration, the mean trail elongation velocity increases linearly, if the number of trails remain constant, a lesser fraction of the total number of monomers are now being transported via the trails. The overall effect is a decrease of the transport rate with increasing basal G-actin concentration. Correspondingly, in the 3-state model the steady state fraction of monomers in the freely diffusing state (*f*_*d*_(*t*)) increases with the increase in concentration if other factors are constant and the net effect is a decrease in the transport rate (figure 8 b, blue dotted line). It should be noted however that usually trail nucleation is expected to increase with increasing G-actin concentration and hence the situation discussed in figure 8 b is only valid when the nucleating factors like formin are the limiting reactants and hence an increase in the G-actin concentration does not lead to higher nucleation rates.

#### 3.3.4. Trail length

In all previous simulations, we have been using the length distribution of actin trails observed in the imaging experiments using GFP:Utr-CH (figure 2). To study the effect of trail length, we use constant trail lengths ranging from 2 *µm* to 20 *µm*.

With increasing trail length, the trail elongation velocities decrease (figure 9(a)) since the trails slow down over their lifetime due to depletion of the G-actin concentration from the axon as monomers are incorporated into the trails as they grow, as discussed in section 3.2. The axonal transport rate however increases with increasing trail length as the monomers are transported over longer distances when they are incorporated in the trails. Monomers incorporated in a trail of length *L*_*t*_ are transported over a mean distance of 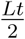. In the 3-state model, even though the effect of decreasing trail elongation velocity is not captured, with increasing trail length there are more monomers in both anterograde and retrograde state, which corresponds to an increase in the steady state fraction of monomers in these states, *f*_*a*_ and *f*_*r*_. The transport velocity hence increases with increasing trail length (figure 9(b), blue dotted line). Experimentally, this can be verified by in-vivo addition of well known actin depolymerization factors like Cytochalasin D, ADF/Cofilin or actin sequestering drugs like Latrunculin A, Thymosin *β*4 [48] which effectively would lead to a decrease in the length of actin filaments thereby leading to a overall decrease in the transport rate.

**Figure 9:**
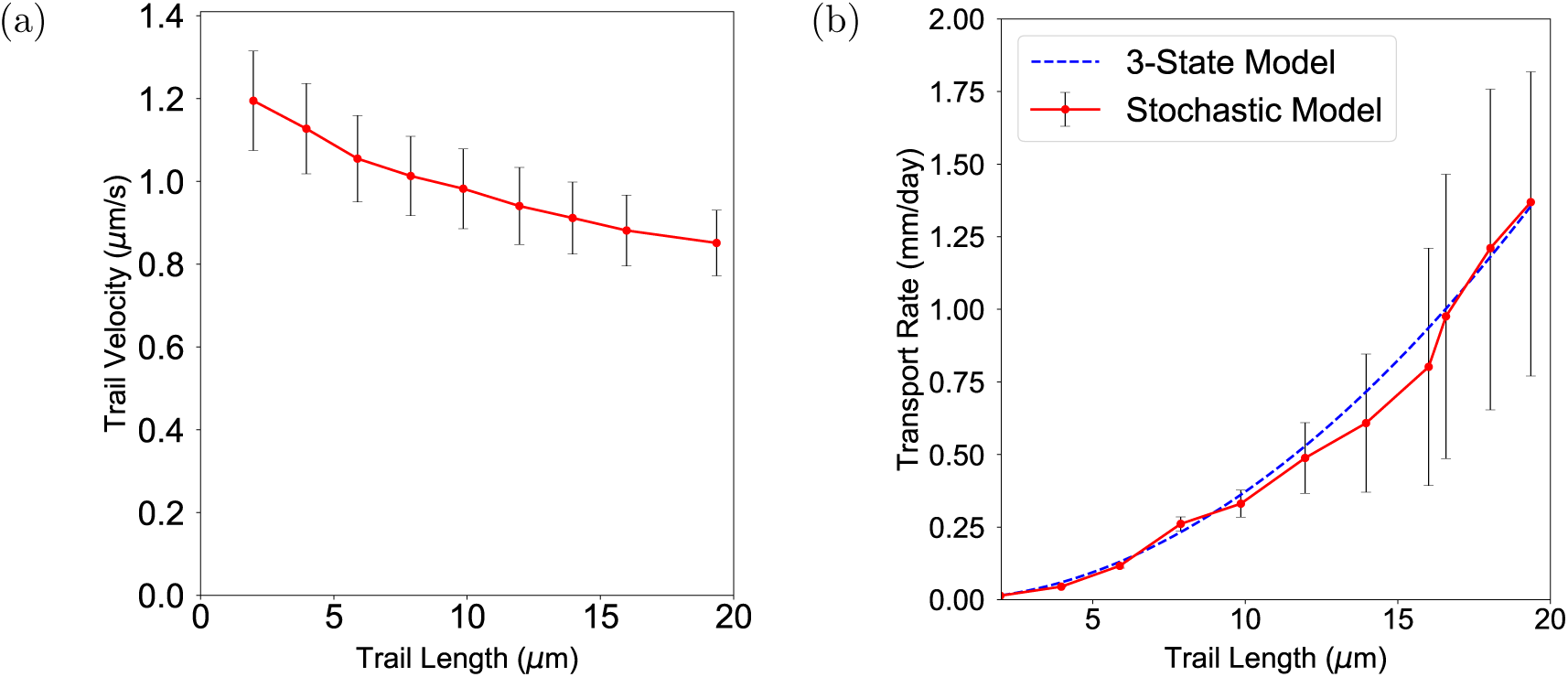
Trail elongation velocity and slow axonal transport rate as a function of the trail length. (a) The trail elongation velocity is averaged over 1000 trails for each length. (b) The transport rate is calcualted for each length by performing a photoactivation simulation and averaging over 1000 runs.

## 4. Conclusions

Does the biased polymerization of actin trails, growing intermittently along the axon, form the mechanistic basis of its active transport? To address this question quantitatively and to identify the major factors that influence this mechanism, we have developed a data-driven stochastic model that incorporates the dynamics of actin trail formation in the axon and a photoactivation simulation method that closely resembles the experimental protocol used to derive in-vivo transport rates of axonal actin. The model is based on actin trails growing linearly along the axon with their barbed ends anchored to stationary hotspots and a small bias in the formation of anterograde trails over retrograde trails. These features of actin trails have been robustly verified by selectively labeling F-actin trails using GFP-Utr:CH and other preferential F-actin markers like Lifeact [23, 21]. Recently, such structures have also been observed in dendrites [49], where filaments have a small anterograde bias (though there is doubt whether they are functionally similar to actin trails) and has been shown to play a role in mediating inter-synaptic vesicle exchange [50].

In an earlier paper [26], we showed that the actin trails can account for the bulk transport of actin along the axon via an active mechanism which does not require the presence of a persistent concentration gradient. This mechanism hinges on the bias of anterograde trails and monomer incorporation at stationary barbed ends. In this paper we discuss how passive diffusion driven by a concentration gradient of G-actin cannot possibly account for the transport rates observed in fluorescence photoactivation experiments. We then discuss the effects of physiological variations in axon and actin trail characteristics on the overall transport rate which also propose new experiments means to further verify this mechanism. We have tested the effect of nucleation rates, axon radius, bulk G-actin concentration and actin trail lengths on actin filament elongation rate and the net transport rate. To better understand how these factors influence the transport rate and to understand why in some cases the transport rate can be inversely related to the filament elongation rate, we have introduced a simplified model where the transition of actin monomers between three distinct kinetic states is considered. This model is analytically solvable and the transport rate can be expressed as a function of the nucleation rate, average trail length, the basal G-actin concentration and the axon radius.

From this model, we find that the mean transport rate is proportional to the trail elongation rate as well as the net fraction of monomers moving in the anterograde direction. The transport rate can be inversely related to the filament elongation rate for increasing axonal radius and G-actin concentration since these factors also decrease the net fraction of monomers moving in the anterograde direction. This model is also helpful in reducing computational complexity since it is analytically solvable. The photoactivation simulations, which we use in the detailed stochastic model, are time consuming and are characterized by a high degree of variability. This variability is mainly due to the fact that photoactivated monomers can bind to trails elongating in both directions and there is only a very small bias of anterograde trails. Averaging over a much higher number of simulations does decrease the variability, but is computationally very costly. The 3-state model helps avoid this since it is analytically solvable and though it makes some simplifying assumptions, the results are qualitatively similar to the results from the detailed stochastic model.

The biased polymerization kinetics of actin is fundamentally different from molecular motor driven polymer sliding models of protein transport and is ideal for cytoskeletal proteins like actin which rapidly exchange between its monomeric and filamentous forms. Though it is unclear how the anterograde bias of trails is generated and maintained persistently, we find that it has profound consequences in regulating bulk transport. Understanding of the biophysics of this bias would help us to further investigate actin transport and how it is disrupted in neurological diseases.

## 5. Acknowledgements

We thank Anthony Brown (Ohio State University, Columbus, OH), Archan Ganguly (University of California, San Diego, La Jolla, CA) and Pankaj Dubey (University of Wisconsin-Madison, WI) for useful discussions and advice. This article is based on research from an ongoing collaboration with Subhojit Roy’s laboratory at the Department of Pathology and Laboratory Medicine and the Department of Neuroscience, University of Wisconsin-Madison, WI.

## Appendix

## Appendix A. Integrating the Diffusion Equation Around the Cross-Section

In section 2.1 we integrated equation 2 around the cross-section since the G-actin concentration is approximately constant around the cross-section of the axon and only considered the one-dimensional diffusion equation. In this section we present results which demonstrate this and outline the detailed integration.

First, we simulate a single actin trail growing in a 3-dimensional axon. The hotspot from which the trail is nucleated is located at (0, 0, 0). The axon is shaped as a parallelopiped with a square cross sectional length of 1.6 *µm*, representing the average radius of the axon and a length of 1000 *µm* stretching from *-*500 *µm* to 500 *µm*. We use periodic boundary conditions along the *x* direction (length of the axon) and reflecting boundary conditions along the *y* and *z* directions (the cross-section). An actin trail is nucleated at *t* = 0 and grows by absorbing actin monomers from the axon. The resultant G-actin concentration along the *x* direction and the *y* direction are plotted in figure A1 Since the G-actin concentration depletion along the radial direction of the axon (figure A1, a) is much less severe compared to the depletion along the length of the axon (figure A1, b), we approximate the concentration to be constant along the radial direction, i.e., *G*(*rr, t*) = *G*(*x, t*). This is expected, since the diameter of the axon (160 nm) is much smaller compared to the characteristic diffusion length of actin. Hence, any concentration change along the radial direction is equilibrated very fast. The details of integrating equating 2 to yield equation 3 are presented below

Starting from equation 2

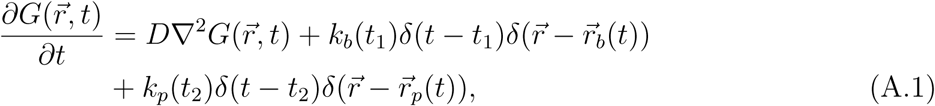

in cylindrical coordinates (*r, θ, x*), if 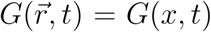 then, 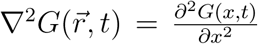. The dirac-delta function in cylindrical coordinates is given by

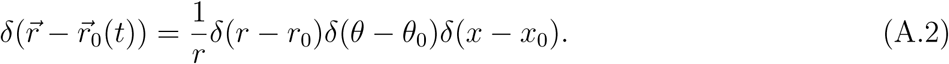

**Figure A1:**
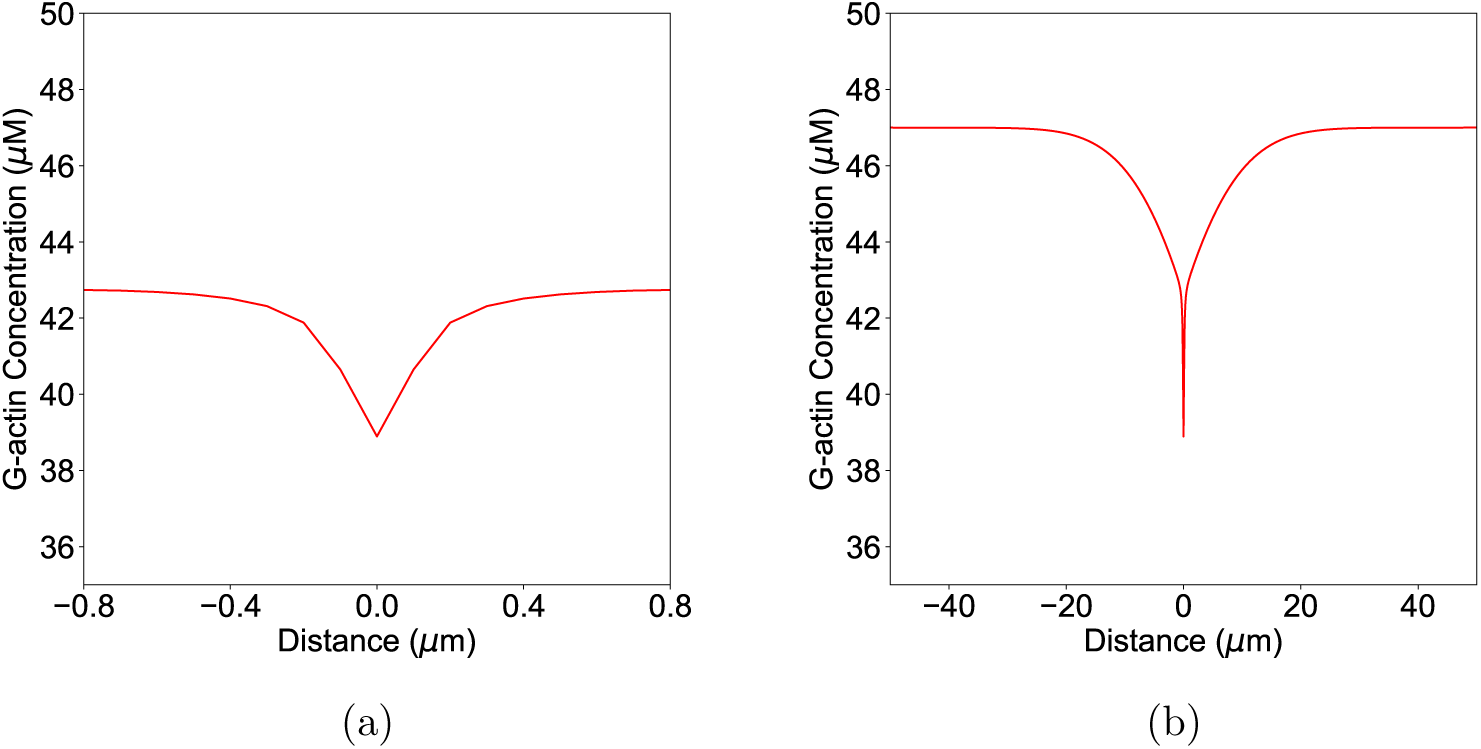
G-actin concentration along the length and radial direction after 7 seconds of trail growth. (a) The G-actin concentration profile in the axon along the radial direction (b) The G-actin concentration profile along the length of the axon.

Since 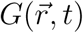 is symmetric in the *r, θ* direction, we project out the *r* and *θ*-integral.

The denominator of equation A.2 becomes

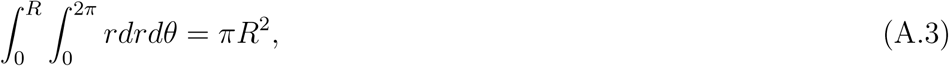

where R is the radius of the axon. Hence, equation A.2 becomes

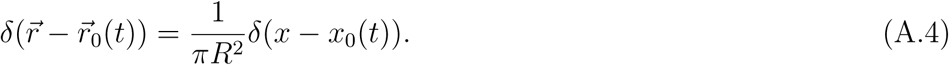

Equation A.1 therefore can be written as

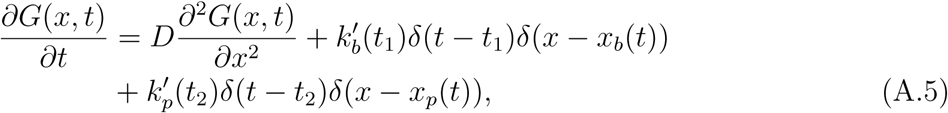

where 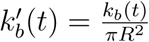 and 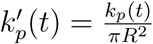 and *R* is the axon radius.

## Appendix B. Quantifying the variability of the transport rate

In the methods section, we have discussed how we determine the transport rate using a photoactivation simulation. In this paradigm, all the monomers in a central zone of the axon are photoactivated. The center of the photoactivated intensity is then tracked over time for hundreds of independent simulation runs. The time averaged center of fluorescence is then fitted to a straight line and the transport rate is defined as the velocity of the center of the photoactivated zone.

**Figure B1:**
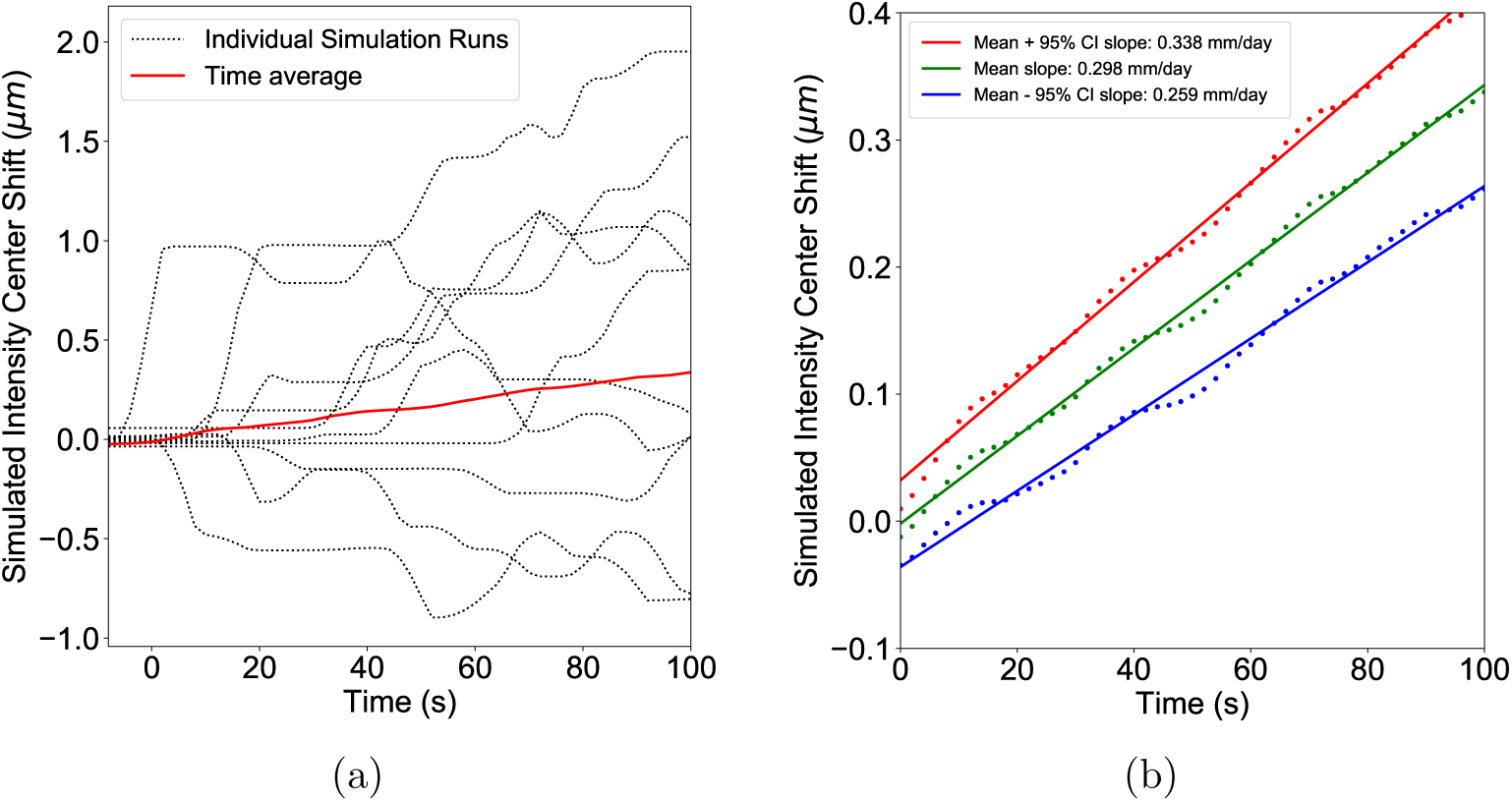
Error estimation for the transport rate calculation. (a) Results from ten individual simulation runs (black, dotted lines) showing the variability of the fluorescence center. The mean (red line) is found by averaging the center of fluorescence over time for 250 independent simulation runs. (b) The transport rate and the error is found by linearly fitting the mean center of fluorescence (blue line) and the 95 % confidence intervals (green and red lines) and determining their slope.

The center of the fluorescence only changes when fluorescently labeled monomers are recruited on an actin trail and individual simulation runs show long periods of pauses interspersed with bouts of movement (figure B1, a, dotted black lines). Therefore, if individually each line is fitted and then the mean of the slopes is characterized as the transport rate, it would be prone to large errors. Instead we take the time average first (figure B1, a, solid red line), which is linear when averaged over many simulation runs and then fit the mean position of the center of fluorescence to a straight line. This procedure is similar to the protocol followed in an in-vivo photoactivation experiment, though the averaging might be over much smaller number of trials (*∽* 50).

Since the variability of the fluorescence position increases with time, a simple variance cannot be defined as the error in this case. Hence, to characterize the variance of this heteroscedastic process, we calculate the 95 % confidence intervals for each time point over all simulation runs. The 95 % confidence intervals are then linearly fitted and their slopes are defined as the error. Figure B1, b shows the mean and the 95 % confidence intervals for 250 simulation runs. The slope of the mean + 95% Confidence Interval line (figure B1, b, green line) is found to be 0.338 mm/day and that of the mean − 95% Confidence Interval line (figure B1, b, red line) is found to be 0.259 mm/day while the mean is 0.298 mm/day (figure B1, b, blue line). The total variance is hence 0.338 mm/day and the assymmetric error bars (0.040 and 0.039 mm/day) are plotted.

